# Representing biological concepts as directions in foundation model embeddings

**DOI:** 10.64898/2026.10.10.758204

**Authors:** Ahmad Shamail, Claire D. McWhite

## Abstract

Biological foundation models encode rich information about protein structure, function, and cellular state in high-dimensional embeddings, but accessing specific biological properties typically relies on additional, task-specific prediction models. Here, we show that diverse biological properties can be mapped to interpretable, reusable directions in embedding space. In the Concept Activation Vector (CAV) approach, examples representing a concept are distinguished from a background set using a linear model, and the resulting weight vector defines the CAV for that concept. In protein embeddings, CAVs recover Gene Ontology and Enzyme Commission annotations, localize Pfam domains, reveal novel structural relationships, and capture DNA-binding and protein-interaction interface specificity. In single-cell embeddings, CAVs isolate cancer-associated expression transitions, and gene variation along these axes substantially agrees with conventional differential-expression analysis despite being a distinct computational approach. Together, these results establish concept directions as a common representation, enabling diverse biological features to be compared, localized, and manipulated.

## INTRODUCTION

Biological foundation models learn high-dimensional representations that encode information about biological structure, properties, and function. Protein language models, for example, learn contextual relationships among amino acids that reflect biochemical and structural patterns, while single-cell foundation models capture patterns of gene expression associated with cellular states. The resulting embeddings support diverse downstream applications. support diverse downstream applications. For proteins, these include structure prediction and design (Madani et al. 2023; Lin et al. 2023), homology search (Kaminski et al. 2023; W. Liu et al. 2024), coevolutionary analysis (Z. Zhang et al. 2024), and a broad range of functional prediction tasks (Leclercq and Droit 2026; J.-Y. Chen et al. 2025). For single cells, foundation models have been applied to cell-type annotation, data integration, gene-network inference, and perturbation-response prediction (Cui et al. 2024; Hao et al. 2024; Baek et al. 2025). However, while these representations are increasingly general-purpose, the methods used to interrogate them remain largely task-specific. Individual biological properties are therefore typically extracted using task-specific prediction models, without explicitly identifying reusable representations of those properties within the embedding space.

This creates a mismatch between the generality of foundation-model representations and the specificity of many biological questions. Researchers may wish to investigate properties outside existing annotation vocabularies, such as a newly identified structural motif, a shared interaction interface, or a transcriptional program associated with a cellular transition. In each case, the relevant biological signal may already be encoded in the pretrained representation, but existing approaches provide limited flexibility for turning that signal into a reusable representation of the biological concept. Moreover, properties such as protein function, enzyme activity, domain identity, interaction specificity, and cellular state are typically addressed by separate prediction systems with different outputs and scoring schemes, making it difficult to compare them directly.

Concept Activation Vectors (CAVs), originally developed for interpreting neural networks (Kim et al. 2018), represent human-interpretable concepts as directions within a model’s internal representation. For example, a CAV can define a direction corresponding to “striped,” and the associated TCAV framework measures how sensitive a model’s prediction of “zebra” is to that concept (**Figure 1A**). Here, we instead use the concept direction itself as the representation of interest, directly measuring how strongly biological embeddings align with it. Given examples such as proteins belonging to a Pfam family (**Figure 1B**) or cells sharing a transcriptional state (**Figure 1C**), we train a linear classifier to distinguish those examples from a background distribution. We then discard the classifier except for its weight vector, which defines the corresponding concept direction. New proteins, sequence regions, residues, or cells can then be scored by their alignment with this vector.

**Figure 1.**
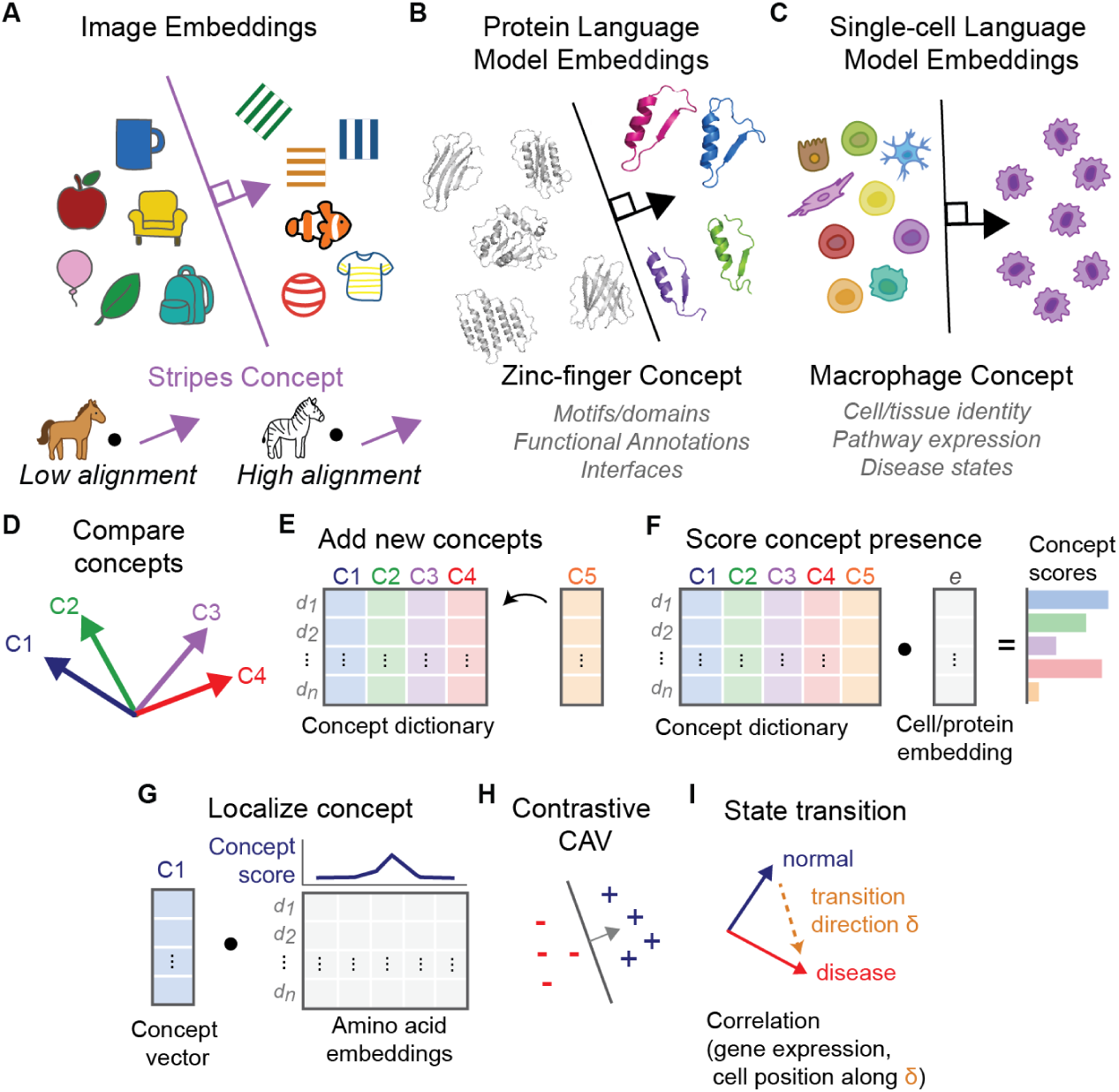
Concept Activation Vectors represent biological concepts as directions in embedding space. **(A)** A concept such as “stripes” can be represented as a direction in embedding space, with higher alignment to an image of a zebra than to a horse. **(B)** A zinc-finger CAV is learned from protein language model embeddings by separating concept examples from a background set. **(C)** The same framework can be applied to single-cell embeddings, illustrated with a macrophage concept. **(D)** CAVs learned in a common embedding space can be directly compared. **(E)** New concepts can be added independently without retraining existing CAVs. **(F)** Protein or cell embeddings can be scored for alignment with each concept by dot product. **(G)** The same operation can be applied to residue embeddings to localize concept-associated signal within a protein. **(H)** Contrastive CAVs can distinguish closely related concepts. **(I)** Differences between concept directions can define transitions between biological states.

The resulting CAV is a reusable representation that can be compared, manipulated, and applied independently. CAVs learned against a common background can be directly compared and clustered (**Figure 1D**), and new concepts can be added independently without retraining existing ones (**Figure 1E**). Protein and cell embeddings can be rapidly scored for alignment with a concept by dot product (**Figure 1F**) and the same alignment operation can be applied to amino acid or sequence-window embeddings to localize concept-associated signal within a protein (**Figure 1G**). Contrastive CAVs can distinguish closely related biological properties (**Figure 1H**). Because the concepts are represented as vectors, standard geometric operations can also be used to define transitions between biological states (**Figure 1I**).

We demonstrate these capabilities across several biological settings. We first ask whether CAVs can recover established protein functional annotations and organize Gene Ontology and Enzyme Commission concepts within a shared space. We then use concept directions to localize protein domains and uncover structural relationships not captured by existing annotations, and show that contrastive CAVs can resolve molecular interaction specificity that is difficult to distinguish using sequence or embedding similarity alone. Finally, we extend the same framework to single-cell embeddings, where differences between concept directions define cancer-associated axes that capture variation in cellular gene-expression states. Together, these analyses test whether biological concepts can be represented as interpretable, reusable directions in pretrained embedding space.

## RESULTS

### CAVs unify and organize protein functional annotations

The function of most proteins is inferred computationally, as relatively few proteins have been experimentally characterized (The UniProt Consortium 2023; Škunca et al. 2017; de Crécy-Lagard et al. 2022).Although homology-based annotation remains highly effective, sequence divergence can obscure relationships among functionally conserved proteins, whereas functional divergence among close homologs can make annotation transfer unreliable. (Schnoes et al. 2009; Rembeza and Engqvist 2021). We therefore asked whether CAVs learned from pretrained protein embeddings could directly recover established functional annotations. Unless otherwise noted, all protein CAVs were trained against the same random protein background, allowing the resulting directions to be compared directly.

We evaluated CAVs across two hierarchical annotation systems: Gene Ontology (GO), which describes molecular functions, biological processes, and cellular components (The Gene Ontology Consortium 2026), and Enzyme Commission (EC) numbers, which classify enzymatic reactions (Tipton and Boyce 2000). Using the proFAB benchmark (Özdilek et al. 2023), comprising 6,678 GO terms and 953 EC terms, we compared CAV-based prediction with two sequence-only deep learning–based annotation methods: DeepGO-SE for GO (Kulmanov et al. 2023) and CLEAN-SupCon for EC (Yu et al. 2023). Both use pretrained protein embeddings to predict predefined functional classes. We additionally included a sequence-similarity baseline using DIAMOND in ultra-sensitive mode (Buchfink et al. 2021; C. Zhang and Freddolino 2024).

For each GO term, we learned a CAV separating annotated proteins from the common background (**Figure 2A**). We then evaluated these CAVs on proteins that received the corresponding GO annotation after the 2021 proFAB dataset cutoff, calculating a log-likelihood ratio (LLR) by comparison to the CAV score distribution of fully withheld term positives and negatives also from the proFAB dataset (**Figure 2B**). We next tested whether the correct GO annotation could be identified among competing terms for each protein. Because highly heterogeneous GO terms may be less well represented by a single linear direction, terms were stratified into diversity bins according to the number of sequence families represented among their annotated proteins (<20, 20–59, or ≥60 families). Candidate terms were ranked by CAV score, with LLR ≤ 0 treated as no prediction. For terms represented by fewer than 60 families, CAVs placed the correct Molecular Function, Biological Process, and Cellular Component annotations among the highest-ranked terms more often than DeepGO-SE or DIAMOND-based sequence similarity (**Figure 2C**). Term-wise evaluation further showed that CAVs achieved high AUROC (**Figure 2D**) and AUPR (**Figure 2E**) across all three GO branches.

**Figure 2.**
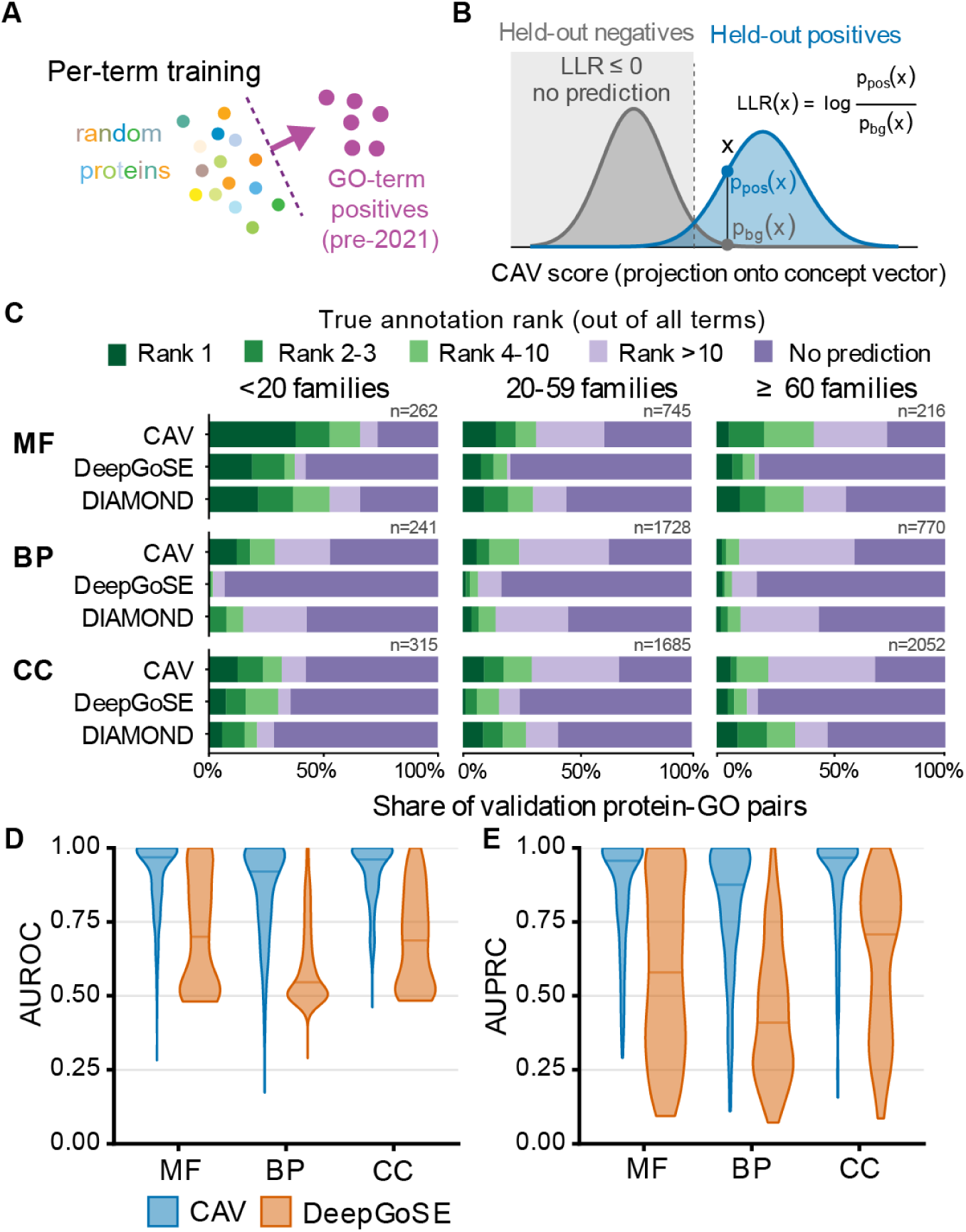
CAVs recover newly annotated Gene Ontology functions. **(A)** For each GO term, a CAV was trained using pre-2021 annotated proteins against a common background of random proteins. **(B)** Log-likelihood ratios (LLRs) are calculated by comparison to withheld negative and positive examples. **(C)** Rank of the true GO annotation among all candidate terms for CAVs, DeepGO-SE, and sequence similarity to training examples (DIAMOND ultra-sensitive) shown separately for Molecular Function (MF), Biological Process (BP), and Cellular Component (CC). Terms are divided by the number of protein families contained within them (<20 families, 818 pairs; 20-59 families, 4,158 pairs; ≥60 families, 3,038 pairs). Predictions with LLR ≤ 0 were treated as no prediction. **(D–E)** Distribution of term-wise AUROC (**D**) and AUPRC (**E**) for CAVs and DeepGO-SE across the three GO branches.

We then turned to EC classification, which organizes enzyme reactions hierarchically from broad classes to sub-subclasses, with complete four-level EC numbers identifying individual reaction categories. For four-level EC classes, a single linear CAV per category performed comparably to DIAMOND in ultra-sensitive mode and CLEAN-SupCon (**Figure 3A**). Performance was weaker at broader EC levels but increased at more specific levels of the hierarchy (**Figure 3B**). This suggests that narrowly defined enzymatic activities are more consistently represented by shared directions in protein embedding space, whereas broader EC classes encompass more mechanistically diverse proteins and may be less well captured by a single linear direction.

**Figure 3.**
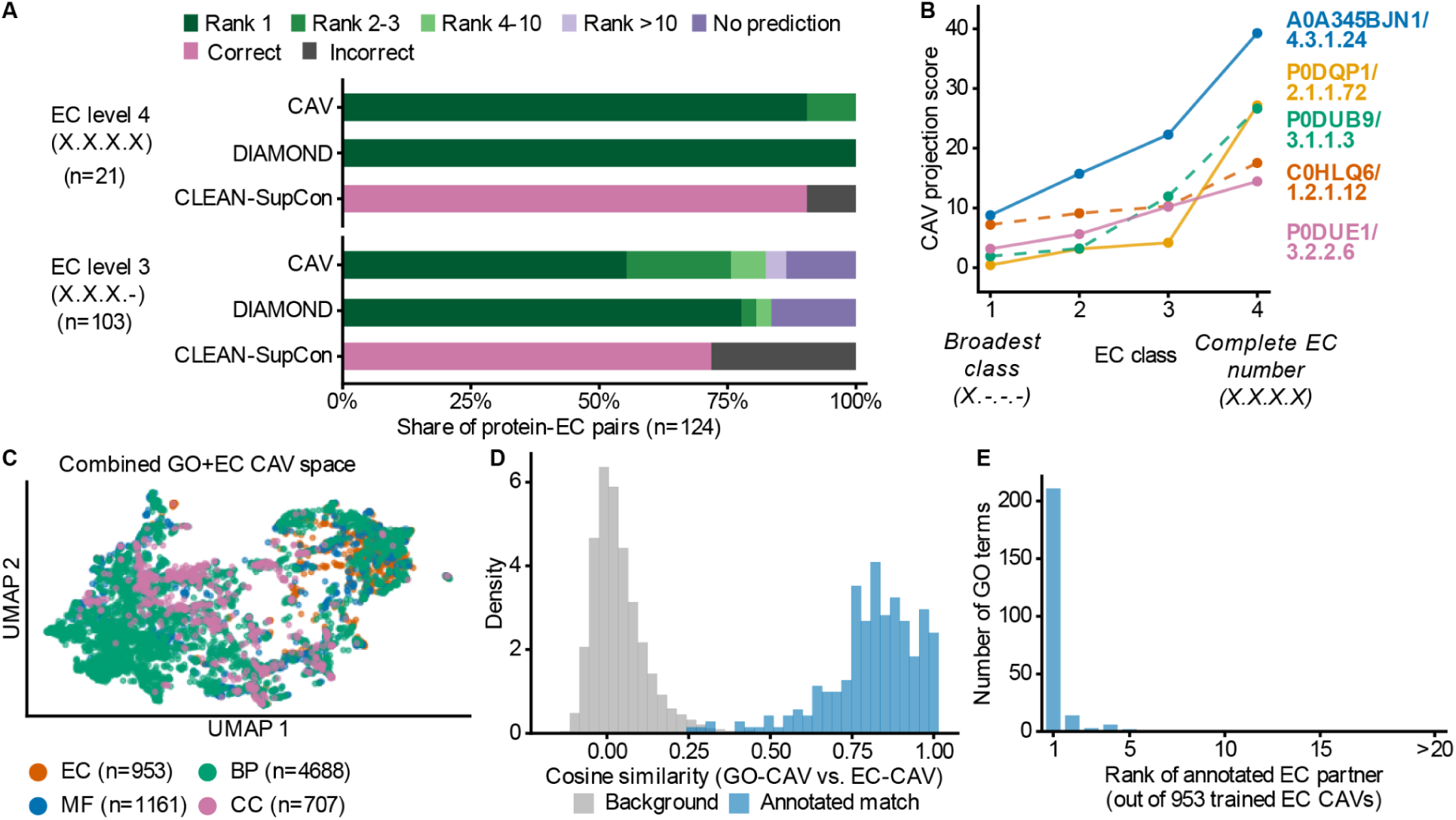
CAVs recover enzyme classifications and relationships between functional ontologies. **(A)** Recovery of EC annotations by CAVs, training-protein sequence similarity (DIAMOND, ultra-sensitive mode), and CLEAN-SupCon for 124 validation protein–EC pairs at EC levels 4 (complete EC number) and 3. CAV predictions are shown by rank of the correct annotation. **(B)** CAV projection scores for representative proteins across successive levels of the EC hierarchy, from broad reaction class to complete EC number. **(C)** UMAP visualization of GO and EC CAVs in their shared embedding space, colored by ontology or GO branch. **(D)** Cosine similarity between GO and EC CAVs for curated EC2GO annotation pairs compared with *unrelated GO–EC pairs. **(E)** Rank of the curated EC partner for 239 GO terms among 953 trained EC CAVs, based on GO–EC CAV cosine similarity*.

A key advantage of representing annotations as CAVs is that concepts from different annotation systems can be compared directly within the same embedding space, as long as CAVs are constructed using a consistent training framework. GO and EC CAVs occupied a shared functional space (**Figure 3C**), and GO–EC pairs linked by the GO Consortium’s curated EC2GO mapping showed substantially higher cosine similarity than unrelated pairs (**Figure 3D**). For 88% of the 239 GO terms represented in both EC2GO and our dataset, the curated EC partner was the most similar of 953 EC CAVs, with most remaining partners also ranked near the top (**Figure 3E**). Thus, the CAV space recovers known relationships between GO and EC annotations without explicitly encoding the mapping between them.

### CAVs distinguish and localize structural motifs

Protein motifs and domains are modular units that encode catalytic, binding, regulatory, and structural functions. Existing resources such as Pfam, InterPro, and PROSITE identify these features largely through sequence profiles, alignments, and curated patterns (Blum et al. 2025; Finn et al. 2014; Sigrist et al. 2026). These approaches are highly effective, but can become less sensitive as sequence similarity declines.

Protein language models provide an alternative representation in which structurally or functionally related regions can remain similar despite substantial sequence divergence (W. Liu et al. 2024; Pantolini et al. 2024). Recent methods have extended motif and domain annotation using sparse autoencoders such as MotifAE, structure-based domain discovery approaches such as TED, and transformer-based sequence segmentation methods such as InterPro-N (Blum et al. 2025; Hou et al. 2025; Lau et al. 2024). Here, we instead ask whether structural motifs can be represented directly as simple directions in pretrained protein embedding space, allowing CAVs to both distinguish motifs and localize them within proteins.

Because CAVs trained against a common background are directly comparable, we hierarchically clustered all Pfam CAVs to generate a nested map of Pfam families (**Figure 4A**). The complete clustering is provided in **Supplemental File 1** and as an interactive visualization at mcwhitelab.org/pfam-circlepack. This organization provides a global view of relationships among Pfam families and a framework for guilt-by-association analysis of poorly characterized domains.

**Figure 4.**
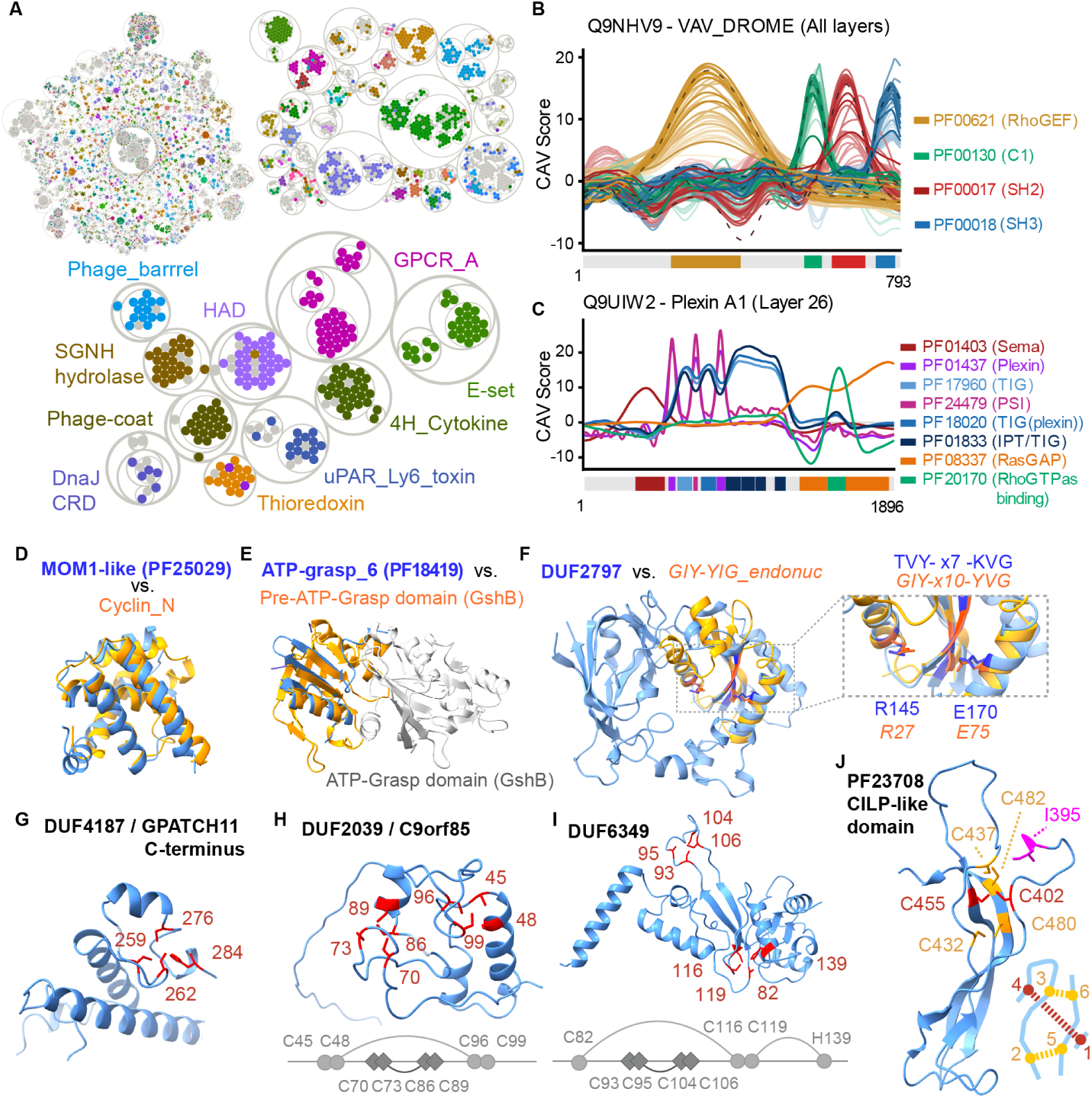
Pfam CAVs reveal global relationships, domain localization, and candidate structural reinterpretations. **(A)** Hierarchical clustering of all Pfam CAVs, visualized as a nested circlepack of Pfam families. Selected clusters are highlighted. **(B)** CAV scores across ESM-C layers for representative domains in VAV_DROME, showing strongest and most localized signal at intermediate layers. **(C)** Domain localization in human Plexin-A1 using layer 26 embeddings; distinct Pfam CAVs peak at their corresponding annotated domains. **(D)** Structural alignment of MOM1-like/PF25029 with a cyclin-like domain. **(E)** Structural alignment of ATP-grasp_6 /PF18419 with the pre-ATP-grasp domain of GshB rather than the ATP-grasp domain. **(F)** Structural comparison identifying a cryptic GIY-YIG-like endonuclease fold within DUF2797/PF10977, with inset highlighting corresponding conserved residues and motif features. **(G)** Candidate Zn-binding site in DUF4187/PF13821 at the C terminus of GPATCH11. **(H)** Two candidate C4 Zn-binding sites in DUF2039/C9orf85/PF10217, including a nested cysteine arrangement. **(I)** Candidate Zn-binding architecture in DUF6349/PF19876, in which a discontinuous C3H site wraps around a compact C4 site. (**J)** Candidate growth-factor-like cystine knot in the CILP-like domain (PF23708) of CILP1, with the 1-4 disulfide threading a macrocycle formed by the 2-5 and 3-6 disulfides. Lumbar disc disease-linked position I395 is highlighted in magenta, with the preceding Ig-like domain omitted for clarity.

We next examined how motif signal varies across model layers. CAV scores were strongest and most sharply localized at intermediate ESM-C layers, particularly layers 21–31, while early and late layers showed weaker signal (**Figure 4B**), consistent with intermediate layers capturing transferable higher-order features (Zeiler and Fergus 2013; Yosinski et al. 2014). Layer 26 was therefore used for subsequent analyses. Using layer 26, we illustrate CAV-based motif localization in the large multidomain human protein Plexin-A1 (**Figure 4C**). CAVs corresponding to different domains produce distinct peaks that closely coincide with their annotated boundaries. For clarity, we show raw CAV scores, which in this example delineate domains at values above approximately 5; these scores can also be calibrated as z-scores against negative examples.

We hypothesized that the structure of the Pfam CAV map could expose relationships and structural features not captured by existing domain annotations. Across the map, we identified numerous candidate reinterpretations of poorly characterized domains, ranging from previously unrecognized fold similarities and putative catalytic domains to unannotated metal-binding architectures. Here, we highlight several examples for which, to our knowledge, the inferred relationships have not previously been proposed in the literature or online domain resources.

As an example of structural inference, we find that PF25029, a domain within the chromatin-associated silencing protein MOM1, adopts a cyclin-like fold (**Figure 4D**). PF25029 is currently defined only by its association with MOM1 and has no assigned molecular function. We also identify potentially misleading Pfam annotations. PF18419, currently annotated as “ATP-Grasp_6/ATP-grasp-like domain,” clusters strongly with pre-ATP-grasp domains. Structural comparison further shows that PF18419 resembles the pre-ATP-grasp domain of GshB rather than its ATP-grasp domain (**Figure 4E**).

We next highlight DUF2797 as a striking example of a cryptic domain embedded within a larger, otherwise uncharacterized domain. Our approach identifies a GIY-YIG-like endonuclease fold hidden within the larger DUF2797 architecture (**Figure 4F**). This relationship was not recovered by Foldseek (van Kempen et al. 2024), illustrating how CAV-based comparisons can complement structure-based searches. This finding provides both a structural reinterpretation of DUF2797 and a specific hypothesis for endonuclease activity.

The Pfam CAV map also revealed a large set of candidate zinc-binding motifs within previously unannotated domains. We identified 84 Pfam families that clustered with known zinc-binding domains and contained structurally plausible candidate Zn-binding sites **(Supplemental File 2).** We highlight several examples spanning distinct architectures. We first identify a candidate CCCH zinc-binding module in the uncharacterized domain PF13821/DUF4187 (**Figure 4G**). This domain frequently occurs at the C-terminus of GPATCH11, a protein implicated in retinal and neurodevelopmental disorders in humans (Zanetti et al. 2024).

We next highlight two candidate zinc-binding domains with less canonical architectures. In DUF2039 (PF10217), the sole annotated domain of the uncharacterized human protein C9orf85, we identify two candidate C4 zinc-binding sites (**Figure 4H**). These sites are organized in a nested AA-BB-BB-AA cysteine arrangement, conceptually similar to the nested RING0 domain of Parkin. Interestingly, C9orf85 was recently found to be physically associated with the BRISC complex (Fischer et al. 2025). As a second example, DUF6349, an uncharacterized domain found largely in proteobacteria, contains an interesting discontinuous C3H site that wraps around a compact C4 site (**Figure 4I**).

Finally, we identify a candidate growth-factor-type cystine knot in the CILP-like domain (PF23708) of cartilage intermediate layer protein 1 (CILP1; UniProt O75339; human residue numbering). This knot architecture appears conserved across chordates, with one disulfide (C402-C455) threading through a nine-residue macrocycle closed by C432–C480 and C437–C482. Interestingly, I395T/rs2073711, a variant associated with intervertebral disc degeneration (J. Liu et al. 2022; Seki et al. 2005), lies adjacent to the predicted knot and is oriented toward it (**Figure 4J**). Notably, the fifth and sixth cysteines required to complete the knot fall outside the current PF23708 domain boundary.

Together, these examples show that CAV-based motif clustering can recover structural relationships that extend beyond existing annotations and generate specific hypotheses for previously uncharacterized domains.

### CAVs can capture binding-interface specificity

As CAVs can be created from arbitrary sequence selections, we next tested whether they could capture binding specificity at molecular interaction interfaces. As a first test, we embedded a single example of the DNA-binding region of human MYC and trained a CAV against our standard background set of proteins (**Figure 5A**). With strong L2 regularization (C = 10⁻⁴), the CAV trained in the full embedding space closely aligned with the original MYC embedding, effectively behaving as a similarity search that preferentially recovered closely related bHLH-Zip proteins. We therefore restricted the embedding space to 128 principal components before training. This shifted the proteome-wide search toward more distantly related bHLH proteins that recognize the same CACGTG motif, including HES/HEY factors. All 20 top-ranked proteins belonged to the bHLH family, and 16 were known CACGTG binders (**Figure 5B**). Of the remaining four, one is a pseudogene, while the others recognize the closely related E-box motifs TACGTG and CAGCTG. Recovering this specificity required pooling embeddings over local amino-acid regions rather than entire proteins, as whole-sequence pooling appeared to dilute the binding-site signal.

**Figure 5.**
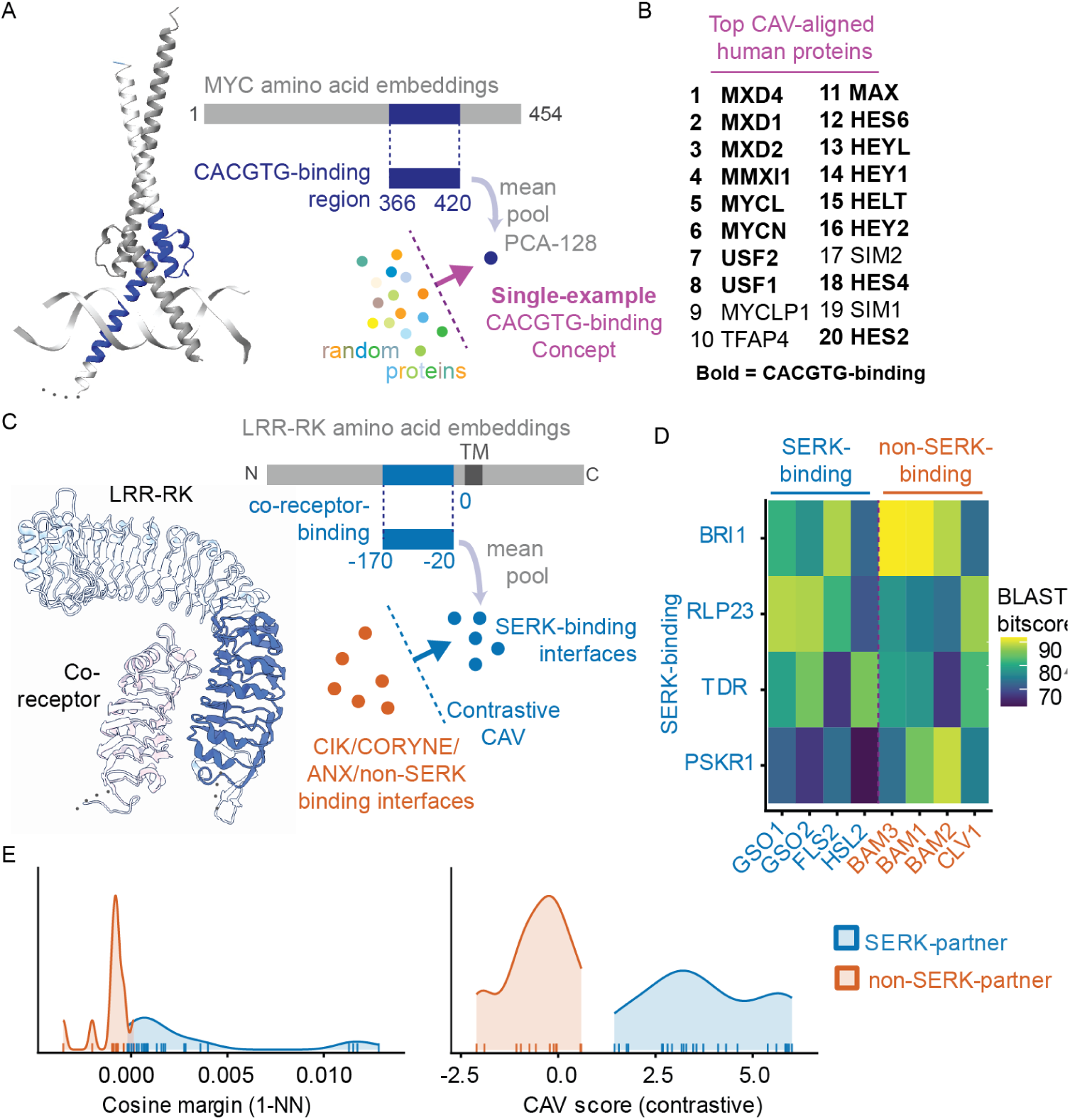
CAVs capture binding-interface specificity from local sequence regions. **(A)** A single-example CAV was constructed from the CACGTG-binding region of human MYC (residues 366–420) by mean-pooling residue embeddings over the selected region and projecting them to 128 principal components. **(B)** Top human proteins aligned to this single-example CACGTG-binding CAV; most are known CACGTG-binding transcription factors. **(C)** A contrastive CAV for LRR-RK co-receptor specificity was trained from membrane-proximal co-receptor-binding regions, using SERK-binding interfaces as positives and CIK/CORYNE/ANX/non-SERK-binding interfaces as negatives. **(D)** Sequence similarity among representative LRR co-receptor-binding regions, showing that SERK-binding specificity is not fully explained by primary-sequence similarity. **(E)** Separation of SERK- and non-SERK-binding receptors by nearest-neighbor cosine margin in the underlying embeddings (left) and by the resulting contrastive CAV score (right).

We then tested the more complex problem of leucine-rich repeat receptor kinase (LRR-RK) co-receptor specificity (**Figure 5C**). Despite the widespread use of shared co-receptors by plant LRR-RKs, determining which receptors preferentially pair with which co-receptors remains a major challenge (Smakowska-Luzan et al. 2018; Xi et al. 2019). For each receptor, we restricted both sequence and embedding comparisons to a membrane-proximal region spanning residues −170 to −20 relative to the transmembrane helix. Although sequence similarity can help predict co-receptor specificity, it does not always distinguish receptors with different co-receptor preferences (**Figure 5D**). For example, the co-receptor-binding region of the SERK-binding receptor BRI1 is more similar in sequence to that of the non-SERK-associated receptor BAM1 than to the corresponding regions of several known SERK-binding receptors.

Nearest-neighbor comparisons using the underlying embeddings were already highly predictive of SERK-binding specificity, indicating that the relevant information was present in the pretrained representation (**Figure 5E**, left). However, the separation occurred over a relatively narrow similarity margin. A CAV trained on SERK-binding interfaces against the common random protein background failed to sharpen this distinction, likely because it captured a more generic co-receptor interface signal. We therefore trained a *contrastive* CAV using 14 SERK-binding *Arabidopsis* LRR interfaces as positives and 18 non-SERK-binding interfaces as negatives. This CAV perfectly separated SERK-from non-SERK-binding receptors in a test set composed entirely of 36 non-*Arabidopsis* proteins (AUROC = 1.00; **Figure 5E, right**). Thus, the contrastive CAV captures a binding-specificity signal already present in the embedding space and represents it as a direction that generalizes across species.

### CAVs organize single-cell gene expression

Thus far, we have used CAVs to identify concepts encoded in protein sequence embeddings. However, the framework is not specific to protein representations and can be applied to other biological feature spaces. We therefore asked whether CAVs could also identify concepts in single-cell gene-expression embeddings, where representations capture cellular expression profiles rather than protein sequences.

We applied CAVs to a multi-tissue single-cell tumor microenvironment atlas containing cells from tumors and adjacent non-tumor tissues (Guimarães et al. 2024). For each cell type in each tissue, we trained two CAVs against a common background of 10,000 randomly sampled cells: one on tumor cells and one on non-tumor cells (**Figure 6A**). We defined the transition direction as the normalized difference between the tumor and non-tumor CAVs. Projecting individual cell embeddings onto this direction provides a continuous score representing their position along the tumor-associated transition axis (**Figure 6B**).

**Figure 6.**
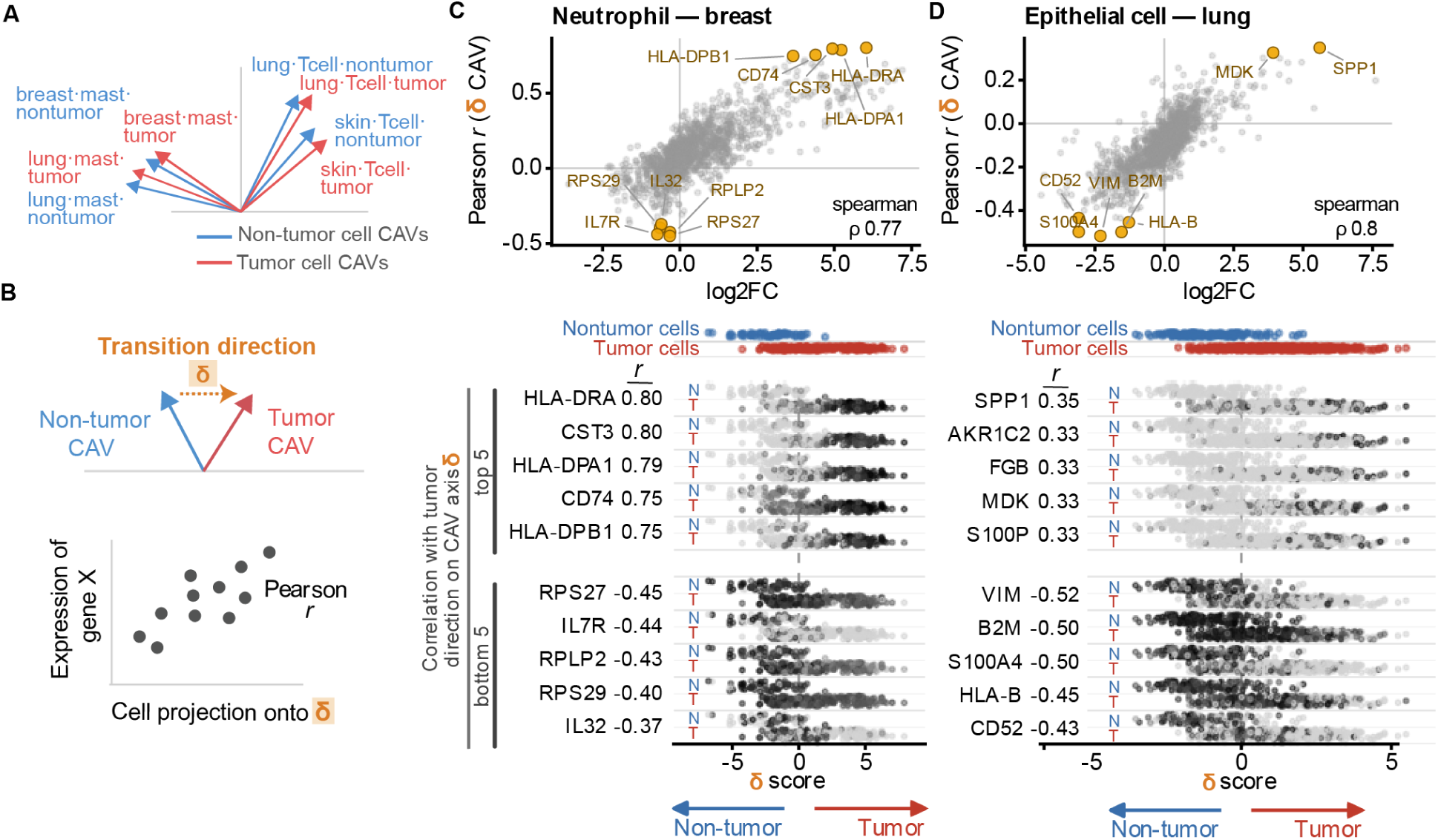
CAVs organize single-cell expression variation into cancer-associated transition axes. **(A)** CAVs can be determined for multiple conditions, and are comparable to each other when trained against a common background **(B)** For each cell type and tissue, the disease axis was the unit difference between the disease CAV and the matched normal CAV. Cells were scored by projecting their embeddings onto this axis. Gene correlates were Pearson correlations between each gene’s log-normalized expression and the cell scores. **(C–D)** Comparison of tumor-versus-non-tumor differential expression (log2FC; x-axis) with the Pearson correlation between gene expression and position along the transition direction axis (y-axis) for breast neutrophils **(C)** and lung epithelial cells **(D).** Selected genes are labeled. Lower panels in C–D show expression of the five most positively and negatively correlated genes across cells ordered by transition direction score, with tumor and non-tumor cells indicated above each strip. Continuum genes that were not tested by pseudobulk DE (removed by the expression filter: AKR1C2, FGB, S100P) appear in the strips but not in the scatter.

We examined two representative transitions spanning distinct cellular contexts: neutrophils in breast cancer (**Figure 6C**) and epithelial cells in lung cancer (**Figure 6D**). We next asked which genes varied along each cancer-associated transition axis and whether these patterns agreed with conventional differential expression. For each case study, we correlated the expression of each gene across individual cells with their position along the transition axis and compared these correlations with normal-versus-cancer differential expression. Differential expression was estimated from donor-level pseudobulk counts using DESeq2. Both analyses were restricted to donors with samples from both normal and cancer conditions.

The two approaches showed substantial agreement for neutrophils in breast cancer (Spearman *ρ* = 0.77) and epithelial cells in lung cancer (*ρ* = 0.80). Strong concordant signals included MHC class II genes (HLA-DRA, HLA-DPA1, and CD74) in neutrophils, consistent with the HLA-DR⁺CD74⁺ antigen-presenting neutrophil state described across human cancers (Wu et al. 2024). In lung epithelial cells, S100P, FGB, and AKR1C2 were among the top CAV-associated genes but were excluded from differential expression testing by standard expression filtering (edgeR::filterByExpr), as each was strongly expressed in tumor samples from only two or three of the four donors. Thus, genes with strong but heterogeneous tumor-associated expression may be excluded by pseudobulk expression filtering while remaining detectable through per-cell correlations with the CAV axis.

CAV-prioritized gene rankings were largely robust to regularization (Spearman ρ = 0.97 for breast neutrophils and 0.91 for lung epithelial cells between C = 1 and C = 10⁻⁴), with 33 and 27 of the top 40 genes shared, respectively. In breast neutrophils, HLA-DRA and HLA-DPA1 ranked first and third at both settings. In lung epithelial cells, genes associated with the tumor end of the axis included keratins (KRT7, KRT18, and KRT19), SPP1, S100P, MDK, and fibrinogen genes, with these associations retained across regularization settings.

These results show that CAV-defined axes capture tumor-associated expression variation consistent with conventional differential expression, while providing a continuous measure of variation across individual cells. Unlike differential expression, which compares expression between conditions, the CAV approach assigns continuous scores to individual cells, allowing expression variation to be examined both across and within conditions.

## DISCUSSION

Representing biological concepts as directions in embedding space provides a simple framework for connecting accumulated biological knowledge with the representational power of biological foundation models.Concept Activation Vectors (CAVs) are fast to train and reusable, allowing proteins to be searched for properties of interest. Because CAVs trained within the same embedding space share a common coordinate system, they can also be compared directly, allowing relationships to be examined within and across annotation systems. In some cases, useful concept directions can be defined from a single example, and their vector representation enables standard operations for biological analysis. The same concept directions can be projected onto local sequence representations to identify where a property is encoded, while contrastive CAVs can isolate specificity between closely related biological properties. CAV-based comparison also generated structural hypotheses for poorly characterized domains, including candidate zinc-binding sites in 84 Pfam families, suggesting a route toward connecting poorly annotated sequences to known structural and functional classes.

In single-cell embeddings, projection along a CAV axis recovered gene-level changes that substantially agreed with conventional differential-expression analysis despite being obtained through a distinct computational approach. By assigning continuous scores to individual cells, CAV-defined axes can also reveal expression variation beyond conventional differential-expression testing, including signals excluded by standard expression filtering.

Despite the simplicity of a linear separator, directions in embedding space can capture protein and gene-expression properties that are often modeled using substantially more complex approaches. This likely reflects the representational work already performed during pretraining: complex sequence or expression relationships are reorganized into a high-dimensional space in which a shared biological property can appear as a coordinated shift across many dimensions. A CAV therefore does not learn these features from scratch, but instead identifies the weighted combination of embedding dimensions that varies consistently with the concept. Even when the underlying features and their interactions are complex and nonlinear, pretraining can therefore transform them into signals that are linearly accessible. Consistent with this interpretation, features often become increasingly linearly separable with depth in neural networks (Alain and Bengio 2016), and structural information can be recovered from protein language-model embeddings using linear projections (Rives et al. 2021). At the same time, predictive performance alone does not uniquely determine the utility of a concept direction for querying or comparison. Multiple directions can separate the training examples equally well yet differ in what they retrieve and in their relationships to other CAVs. In particular, PCA dimensionality, feature scaling, and regularization strength can substantially alter the learned direction, potentially changing which proteins are retrieved and how concepts relate to one another, even when classification performance remains similar.

CAVs have been applied to biological and biomedical models since their introduction, most often in the context of concept attribution. The original TCAV study used retinal lesions to interpret predictions from a diabetic-retinopathy classifier (Kim et al. 2018), and subsequent work has used concept vectors to relate histological features to model predictions (Graziani et al. 2020; Gamble et al. 2021) and to identify and remove unwanted demographic signals in medical-imaging models (Correa et al. 2024). More recently, TPCAV extended concept attribution to genomics, enabling regulatory motifs and chromatin features to be related to deep-learning predictions (Yang and Mahony 2026). These studies establish concept attribution as a useful way to connect biologically meaningful features to model behavior.

Concept directions have also begun to support uses beyond attribution. Text2Concept derives CAVs from natural-language descriptions and uses similarity to these directions to rank images and perform zero-shot classification (Moayeri et al. 2023). CAVs have likewise been used to modify generated 3D shapes by adding or combining concept-associated directions in latent space (Druc et al. 2022) and to steer large language models by adding or removing learned concept directions during inference (H. Zhang et al. 2025).

Like conventional linear probes, CAVs use linear classifiers to identify properties encoded in pretrained representations. However, whereas linear probing typically focuses on predictive performance, we treat the learned directions themselves as reusable representations of biological concepts that can be compared, localized, contrasted, and used to search for related examples. This framework could complement task-specific prediction systems by providing a lightweight way to define and reuse biological properties within a common representation. Once embeddings have been generated, new concepts are inexpensive to define and can be added independently without retraining the underlying foundation model or existing CAVs. While embedding generation is generally most efficient on GPUs, CAV training and scoring are lightweight and can readily be performed on CPUs.

At the same time, CAVs can recover only information that is present in the underlying representation, and their performance will inherit the limitations, biases, and model-specific properties of that embedding space. Concept signal should therefore be established independently for each application rather than assumed to be present. The property captured by a CAV also depends on the background or contrast used to define it, allowing related signals to be progressively refined but making the choice of reference set an important part of concept definition. More broadly, our results suggest a practical workflow for progressively refining CAV-based searches. An initial CAV trained against a generic background can identify candidates that share signal with the concept of interest. Known false positives from this search can then be incorporated as hard negatives in a contrastive CAV, refining the direction toward the more specific biological property that distinguishes true positives from closely related alternatives.

The applications explored here likely represent only a small subset of what can be captured as reusable directions in biological embedding spaces. Because the framework requires only vector representations, it can in principle be applied to many other biological data types and models. As foundation models increasingly integrate sequence, structure, expression, imaging, and other modalities into shared representations, CAVs could provide a common way to define and relate concepts across distinct forms of biological information. The broader opportunity is therefore not any single downstream application, but an extensible vocabulary of biological concepts that can grow with new models, datasets, and biological questions.

## ONLINE METHODS

### Data and code availability

Code for training, analyzing, and applying Concept Activation Vectors is available at https://github.com/mcwhitelab/bioCAV. Code for reproducing paper analyses is available at https://github.com/mcwhitelab/bioCAV-paper. Associated data files, including trained Pfam, GO, and EC CAVs are available at zenodo.org with identifier 10.5281/zenodo.23284980.

### Training concept activation vectors

Biological concepts were represented as concept activation vectors (CAVs) learned from embeddings generated by pretrained language models. All protein sequences were downloaded through the UniProt REST API (Ahmad et al. 2025), and protein sequences were embedded using the ESMPlusPlus_large implementation of ESM-C 600M (Synthyra 2024; ESM Team 2024). Embeddings from layer 26 of the 36-layer model were mean-pooled over the sequence or selected region to obtain a single embedding per input. For single-cell analyses, cells were represented using 768-dimensional embeddings from the penultimate layer of Geneformer-V2-104M(H. Chen et al. 2026). Raw UMI counts were mapped to the Geneformer gene vocabulary, normalized to 10,000 counts per cell, and scaled by gene-specific median expression values from the pretraining corpus. Genes were ranked by their scaled expression, and the top 4,094 were retained. Cell embeddings were extracted from the <cls> token representation.

For each concept, positive examples were defined by the corresponding biological annotation or user-specified example set. Unless otherwise stated, CAVs were trained against a common background of 10,000 randomly sampled proteins from UniRef50 (Suzek et al. 2015) or 10,000 random cells from the CELLxGENE census (CZI Cell Science Program et al. 2025). For each modality, a StandardScaler was fit to the shared background population and used for both CAV training and subsequent scoring. CAVs describing SERK-binding specificity were an exception to the common-background formulation and were trained contrastively, using SERK-binding interfaces as positive examples and non-SERK-binding interfaces as the negative class, with a StandardScaler fit to training examples.

For analyses incorporating dimensionality reduction, PCA was fit to the standardized shared background population, and embeddings were projected onto the top 128 principal components before CAV training. The resulting coefficient vector was then mapped back into the full standardized embedding space, allowing CAVs trained at different PCA dimensions to be scored and compared in the original coordinate frame.

A linear L2-regularized logistic regression (L-BFGS solver, class_weight = “balanced”) was trained independently for each concept to distinguish positive from negative embeddings. Regularization was adjusted in individual analyses to balance recovery of shared concept signatures with sensitivity to within-concept variation, while accounting for training-set size. The fitted coefficient vector, which is orthogonal to the learned decision boundary, was oriented toward the positive class and normalized to unit L2 norm to define the CAV, and the fitted classifier was discarded. Repeating this procedure for each concept produced a dictionary containing one vector per biological concept; addition of a new concept therefore required training only its corresponding CAV and did not require retraining existing vectors.

### CAV scoring

CAVs were queried by measuring the alignment between a query embedding and the corresponding concept vector. Query embeddings were first transformed using the same StandardScaler used during CAV training. For a standardized query embedding (h) and unit CAV (v), the CAV score (s) was calculated as their dot product, ( ^T^ ). Larger scores indicate greater alignment with the direction associated with the target concept. The same scoring operation was used irrespective of the biological input, with query embeddings representing complete proteins, protein subsequences, or individual cells.

### CAV localization and querying

For localized protein analyses, the full-length protein was embedded to obtain residue-level representations. Overlapping sequence windows of the specified length were mean-pooled from these representations at a stride of one residue and scored against the corresponding CAV. To generate a residue-level profile, the score at each sequence position was defined as the mean CAV score across all windows spanning that residue, yielding a continuous measure of concept alignment along the protein. For proteome-wide querying, the highest-scoring window within each protein was used as its match score. For concepts trained on whole-protein embeddings, including GO and EC CAVs, the mean-pooled embedding of the complete protein can be scored directly against the corresponding CAV.

### Gene Ontology and Enzyme Commission Datasets and Evaluation

GO and EC CAV libraries were constructed from the proFAB benchmark datasets (Özdilek et al. 2023). For each CAV, we subsampled the positive random-split train set files to 1000 sequences, and trained against the common protein background of 10,000 sequences using balanced L2-regularized logistic regression with C = 10⁻⁴ in the shared scaler frame. We used C = 10⁻⁴ to favor the dominant contrast between concept-positive proteins and the shared background and emphasize the concept’s common signature. This produced libraries of 6,678 GO CAVs and 953 EC CAVs. To produce per-term LLRs, we produced positive and negative background CAV-score distributions by scoring each CAV against positive and negative random-split test set proteins, subsampling each to 1000 sequences. As the proFAB dataset does not use annotations added after 2020, we computed log-likelihood ratios (LLRs) on a new temporal validation set of protein-GO term pairs newly annotated between 2021-12-31 and 2026-05-27 and evidence code IDA and its descendants, downloaded from QuickGO (Binns et al. 2009).

For EC temporal validation, we used the test_ec.csv file produced by EC-Bench (Davoudi et al. 2026), constructed from protein-EC pairs present in the 2023-01 release of UniProt, but not in the 2018-02 release. We then removed any protein-EC pairs present in the proFAB positive random-split training set. LLRs were computed as for GO terms above.

To assess confidence in CAV-based predictions, we fitted separate Gaussian distributions to the CAV projection scores of held-out positive and negative examples from proFAB. For a query with projection score (s), we calculated the log-likelihood ratio: *LLR*(*s*) = *log*[*p*(*s*|*positive*) / *p*(*s*|*negative*)] where *p*(*s*|*positive*) and *p*(*s*|*negative*) are the Gaussian probability densities of the positive and negative score distributions, respectively. Positive LLR values indicate that a query score is more likely under the positive distribution than under the negative distribution, and predictions with LLR ≤ 0 were treated as no prediction.

For comparison to a state-of-the-art sequence-only GO term prediction model, we ran DeepGO-SE locally with default parameters and threshold of 0.1 for GO terms, and directly used the clean-supcon predictions on the test_ec.csv proteins produced by EC-Bench (data/results_test_100_go_3.csv).

We also included a sequence-similarity baseline using DIAMOND (v2.0.9) (Buchfink et al. 2021) in ultra-sensitive mode with an E-value threshold of 1 and self-hits excluded, as recommended for sequence-based annotation transfer in (C. Zhang and Freddolino 2024). Each candidate GO or EC term was assigned the highest bitscore among hits to its CAV training-positive proteins. Terms were ranked over the same filtered candidate sets used for CAV evaluation, with tied scores assigned their average rank. A validation pair was classified as no prediction when no qualifying hit was found to the true term’s training-positive proteins.

To assess annotation prioritization, each validation protein–term pair was ranked against all trained CAVs in the same GO ontology or at the same EC level. For GO, candidate sets excluded obsolete terms, the true term’s ancestors, and the protein’s other known annotations from QuickGO and proFAB, including their ancestors. For EC, the protein’s other known EC-Bench and proFAB-positive annotations at the same level were excluded. CAVs were ranked by raw projection score. DeepGO-SE predictions were ranked over the same GO candidates, assigning zero to unreported scores. Rank was defined as one plus the number of candidates scoring strictly above the true annotation. Pairs with CAV LLR ≤ 0 or an unreported DeepGO-SE true-term prediction were classified as no prediction. CLEAN-SupCon predictions were evaluated as correct or incorrect rather than ranked. We additionally determined approximate number of protein family per GO term by running DIAMOND all-by-all search on all term members, filtering for edges over 30% identity, and counting connected components.

For GO, AUROC and average precision were calculated separately for each term using identical positive and negative protein sets for CAVs and DeepGO-SE. Positives were the temporal validation proteins remaining after exclusion of the term’s proFAB positive training and test examples. Negatives were the corresponding proFAB random-split negative test proteins, not the shared 10,000-protein training background. The evaluated sets contained 9–77 negatives per term. CAV evaluation used raw projection scores, while DeepGO-SE evaluation used its reported scores, assigning zero to predictions omitted by its 0.1 output threshold. AUROC and AUPRC were computed using scikit-learn’s roc_auc_score and average_precision_score, respectively. Each term contributed one value; scores were not pooled across terms. The reference prevalence for average precision was each term’s positive fraction.

### PFAM Datasets and Evaluation

CAVs were trained for 27,472 Pfam families using sequence regions corresponding to up to 1000 annotated Pfam instances as positive examples. We obtained domain coordinates through the InterPro REST API (Blum et al. 2025). For each positive protein, the full protein sequence was embedded with ESM-C and residue-level embeddings within the annotated Pfam span were mean-pooled to obtain a single representation for that instance. Pfam CAVs were trained against the common background of 10,000 UniRef50 proteins described above. To visualize how localization varied across model depth, CAVs for four Pfam domains in VAV_DROME (PF00621, PF00130, PF00017, and PF00018) were additionally trained at all 36 model layers and their localization profiles compared across layers.

Training the complete library of 27,472 Pfam CAVs from precomputed embeddings required 21 min wall-clock time (45 CPU core-hours; AMD EPYC 7642 CPUs), with a median fitting time of 1.3 s per CAV. These timings exclude generation of the protein embeddings, which were computed once and reused across CAV training.

To localize Pfam concepts within query proteins, full-length proteins were embedded to obtain residue-level representations. For each Pfam family, the scanning-window length was set to the median length of its annotated training instances. Overlapping windows were mean-pooled at a stride of one residue and scored against the corresponding CAV. The score assigned to each residue was calculated as the mean score of all windows spanning that position, producing a continuous localization profile along the protein.

### Pfam clustering

To organise the Pfam CAV library, 27,472 CAVs were clustered using a cosine-similarity nearest-neighbour graph (mutual *k* = 15, minimum similarity = 0.10). Diffusion-map coordinates approximating random-walk distances (*t* = 16) were used for hierarchical clustering with average linkage. Tree heights were normalised to the largest connected component and cut at 0.45, 0.20, 0.10 and 0.05 to define four levels. The hierarchy was visualised as a circle pack using ggraph (Pedersen 2025), with families coloured by Pfam 38.2 clan. Cluster assignments at 20 thresholds (0.05–1.00) are provided in **Supplemental File 1**.

### Zn-finger candidate screen

Candidate Zn-finger and Zn-ribbon containing Pfam families were screened for Cys/His metal-binding sites in AlphaFold2 structures downloaded from Alphafold DB, restricted to InterPro-defined domain boundaries. Up to three proteins per family were examined, and four-residue Cys/His sites were identified using ligand-atom geometry (mean distance to centroid ≤3.0 Å; pairwise distances 2.6–5.0 Å). A family was retained if a site recurred in at least two proteins. The same procedure detected 58/60 known zinc-module families and 0/148 random Pfam controls. Candidates are provided in **Supplemental File 2**.

### Interaction Interface analysis

For the MYC analysis, a CAV was trained using a single annotated MYC DNA-binding region (MYC_HUMAN, residues 366-420), against the common protein background. We used C = 10⁻⁴ to favor the dominant concept-versus-background contrast and fitted the CAV after projecting standardized embeddings onto 128 principal components learned from the background. The resulting CAV was then used to search the human proteome. Each query protein was embedded once to obtain residue-level embeddings, then these embeddings were pooled across spans as described in the CAV Localization section above.

To test whether CAVs could distinguish interaction specificity between closely related receptor interfaces, we trained a contrastive CAV using the approximate co-receptor-binding region of 32 *Arabidopsis thaliana* LRR receptors. For each receptor, we used a 150-residue region ending 20 residues before the transmembrane helix. The positive class consisted of 14 receptors that function with SERK co-receptors, while the 18 hard negatives are comprised of six CIK-dependent receptors (CLV1, CLV2, RPK2, BAM1–3) and 12 SERK-independent receptors (PRK1–6 and SRF2, SRF4–8). Generalization was evaluated on 36 receptors from non-*Arabidopsis* plants, including 24 SERK-binding and 12 non-SERK-binding. Eight receptors were curated independently from rice, tomato, maize, and canola, and the remaining 28 were identified by homology to receptors in the training set. We used balanced L2-regularized logistic regression with C = 1, as the stronger regularization used for the large protein libraries excessively suppressed the discriminating signal in this small training set.

As an embedding-similarity baseline, each test interface was compared with the training interfaces using cosine similarity. A signed nearest-neighbor margin was calculated as the cosine similarity to the closest SERK-binding training receptor minus the similarity to the closest non-SERK-binding receptor, such that positive values indicated greater similarity to the SERK-binding class. BLASTp comparisons of the 150-amino-acid interface regions were additionally used to examine selected receptors for which the nearest homologs from the two interaction classes were closely matched.

### Single Cell Analysis

Single-cell analyses were performed using the CELLxGENE collection “A multi-tissue single-cell tumor microenvironment atlas” (3f7c572c-cd73-4b51-a313-207c7f20f188) (Guimarães et al. 2024). Single-cell CAVs were constructed from combinations of cell type, tissue, and disease state within this dataset. Groups containing fewer than 50 cells were excluded, and up to 500 cells were used as positive examples for each concept. Blood, PBMC, and lymph-node samples were excluded from the CAV library. Cell embeddings were standardized using a shared StandardScaler and projected onto 128 principal components before CAV training. Each concept was trained against the common background of 10,000 randomly sampled CELLxGENE Census cells described above, producing 93 single-cell CAVs. We tested both *C* = 10⁻⁴ and *C* = 1 for the single-cell CAVs as a sensitivity comparison, and final analyses use *C* = 1.

To capture the tumor-associated transition within each cell type, we trained two CAVs for each cell type in each tissue: one on tumor cells and one on non-tumor (tumor-adjacent) cells, both against the common background of 10,000 reference cells. The transition axis was defined as the difference between the two unit-normalized CAV directions, d = v^_tumor − v^_non-tumor, rescaled to unit length. Because both CAVs share the same cell type, tissue and background, components common to the two cancel in the difference, and d points from the non-tumor toward the tumor state of that cell type. Each cell was assigned a continuous transition score by projecting its embedding onto d. Higher scores indicate a more tumor-like state.

To identify genes associated with the transition axis, we restricted the analysis to cells from donors who contributed both tumor and non-tumor cells of the given cell type. Counts were normalized to 10,000 per cell and log1p-transformed, and we computed the Pearson correlation between each gene’s expression and the transition score across cells. We compared gene-level correlations with pseudobulk DESeq2 log2 fold changes using Spearman correlation over the genes tested in both analyses.

For conventional differential-expression analysis, raw UMI counts were summed per donor and condition, using donors represented in both normal and cancer groups. Pseudobulk samples containing fewer than 10 cells were excluded together with the donor’s paired sample. Genes were filtered using edgeR::filterByExpr (Robinson et al. 2010) with default settings. Differential expression was tested using DESeq2 (v1.46.0) (Love et al. 2014) with the design ∼ donor + condition, normal as the reference condition, and default size-factor normalization. We report unshrunk maximum-likelihood log₂ fold changes for cancer relative to normal. Wald-test P values were adjusted using the Benjamini–Hochberg procedure within each comparison. Genes with adjusted P < 0.05 and |log₂ fold change| ≥ 1 were considered differentially expressed.

Both the CAV-based gene-association and differential-expression analyses were restricted to donors represented in both normal and cancer conditions. Three cell-type/tissue combinations were highlighted in the main figure: breast neutrophils, lung epithelial cells, and colorectal fibroblasts; lung epithelial cells were additionally restricted to the 10x 3′ v2 assay. Agreement between CAV-derived gene associations and differential-expression effect sizes was assessed using Spearman correlation across genes.

Supplemental File 1: Clustering of Pfam CAVs

Supplemental File 2: Candidate zinc-binding Pfam domains

## Reference

Ahmad, Shadab, Leonardo Jose da Costa Gonzales, Emily H. Bowler-Barnett, et al. 2025. “The UniProt Website API: Facilitating Programmatic Access to Protein Knowledge.” Nucleic Acids Research 53 (W1): W547–53.

Alain, Guillaume, and Yoshua Bengio. 2016. “Understanding Intermediate Layers Using Linear Classifier Probes.” In arXiv [Stat.ML]. October 5. arXiv. 10.48550/arXiv.1610.01644.

Baek, Seungbyn, Kyungwoo Song, and Insuk Lee. 2025. “Single-Cell Foundation Models: Bringing Artificial Intelligence into Cell Biology.” Experimental & Molecular Medicine 57 (10): 2169–81.

Binns, David, Emily Dimmer, Rachael Huntley, Daniel Barrell, Claire O’Donovan, and Rolf Apweiler. 2009. “QuickGO: A Web-Based Tool for Gene Ontology Searching.” Bioinformatics (Oxford, England) 25 (22): 3045–46.

Blum, Matthias, Antonina Andreeva, Luis Carlos Florentino, et al. 2025. “InterPro: The Protein Sequence Classification Resource in 2025.” Nucleic Acids Research 53 (D1): D444–56.

Buchfink, Benjamin, Klaus Reuter, and Hajk-Georg Drost. 2021. “Sensitive Protein Alignments at Tree-of-Life Scale Using DIAMOND.” Nature Methods 18 (4): 366–68.

Chen, Han, Madhavan S. Venkatesh, Javier Gόmez Ortega, et al. 2026. “Scaling and Quantization of Large-Scale Foundation Model Enables Resource-Efficient Predictions in Network Biology.” Nature Computational Science 6 (5): 450–63.

Chen, Jia-Ying, Jing-Fu Wang, Yue Hu, Xin-Hui Li, Yu-Rong Qian, and Chao-Lin Song. 2025. “Evaluating the Advancements in Protein Language Models for Encoding Strategies in Protein Function Prediction: A Comprehensive Review.” Frontiers in Bioengineering and Biotechnology 13: 1506508.

Correa, Ramon, Khushbu Pahwa, Bhavik Patel, Celine M. Vachon, Judy W. Gichoya, and Imon Banerjee. 2024. “Efficient Adversarial Debiasing with Concept Activation Vector - Medical Image Case-Studies.” Journal of Biomedical Informatics 149 (104548): 104548.

Crécy-Lagard, Valérie de, Rocio Amorin de Hegedus, Cecilia Arighi, et al. 2022. “A Roadmap for the Functional Annotation of Protein Families: A Community Perspective.” Database 2022: baac062.

Cui, Haotian, Chloe Wang, Hassaan Maan, et al. 2024. “scGPT: Toward Building a Foundation Model for Single-Cell Multi-Omics Using Generative AI.” Nature Methods 21: 1470–80.

CZI Cell Science Program, Shibla Abdulla, Brian Aevermann, et al. 2025. “CZ CELLxGENE Discover: A Single-Cell Data Platform for Scalable Exploration, Analysis and Modeling of Aggregated Data.” Nucleic Acids Research 53 (D1): D886–900.

Davoudi, Saeedeh, Christopher S. Henry, Christopher S. Miller, and Farnoush Banaei-Kashani. 2026. “EC-Bench: A Benchmark for Enzyme Commission Number Prediction.” Bioinformatics Advances 6 (1): vbag004.

Druc, Stefan, Aditya Balu, Peter Wooldridge, Adarsh Krishnamurthy, and Soumik Sarkar. 2022. “Concept Activation Vectors for Generating User-Defined 3D Shapes.” In arXiv [Cs.CV]. April 29. arXiv. 10.48550/arXiv.2205.02102.

ESM Team. 2024. “ESM Cambrian: Revealing the Mysteries of Proteins with Unsupervised Learning.” Preprint, EvolutionaryScale Website. https://evolutionaryscale.ai/blog/esm-cambrian.

Finn, Robert D., Alex Bateman, Jody Clements, et al. 2014. “Pfam: The Protein Families Database.” Nucleic Acids Research 42 (Database issue): D222–30.

Fischer, Samantha N., Erin R. Claussen, Savvas Kourtis, et al. 2025. “Hu.MAP3.0: Atlas of Human Protein Complexes by Integration of >25,000 Proteomic Experiments.” Molecular Systems Biology 21 (7): 911–43.

Gamble, Paul, Ronnachai Jaroensri, Hongwu Wang, et al. 2021. “Determining Breast Cancer Biomarker Status and Associated Morphological Features Using Deep Learning.” Communications Medicine 1 (1): 14.

Graziani, M., V. Andrearczyk, S. Marchand-Maillet, and H. Müller. 2020. “Concept Attribution: Explaining CNN Decisions to Physicians.” Computers in Biology and Medicine 123 (103865): 103865.

Guimarães, Gabriela Rapozo, Giovanna Resk Maklouf, Cristiane Esteves Teixeira, et al. 2024. “Single-Cell Resolution Characterization of Myeloid-Derived Cell States with Implication in Cancer Outcome.” Nature Communications 15 (1): 5694.

Hao, Minsheng, Jing Gong, Xin Zeng, et al. 2024. “Large-Scale Foundation Model on Single-Cell Transcriptomics.” Nature Methods 21: 1481–91.

Hou, Chao, Di Liu, and Yufeng Shen. 2025. “MotifAE Reveals Functional Motifs from Protein Language Model: Unsupervised Discovery and Interpretability Analysis.” bioRxiv: The Preprint Server for Biology, ahead of print. 10.1101/2025.11.04.686576.

Kaminski, Kamil, Jan Ludwiczak, Kamil Pawlicki, Vikram Alva, and Stanislaw Dunin-Horkawicz. 2023. “pLM-BLAST: Distant Homology Detection Based on Direct Comparison of Sequence Representations from Protein Language Models.” Bioinformatics (Oxford, England) 39 (10): btad579.

Kempen, Michel van, Stephanie S. Kim, Charlotte Tumescheit, et al. 2024. “Fast and Accurate Protein Structure Search with Foldseek.” Nature Biotechnology 42 (2): 243–46.

Kim, Been, Martin Wattenberg, Justin Gilmer, et al. 2018. “Interpretability Beyond Feature Attribution: Quantitative Testing with Concept Activation Vectors (TCAV).” Preprint. https://arxiv.org/abs/1711.11279.

Kulmanov, Maxat, Francisco Guzmán-Vega, Paula Duek, Lydie Lane, Stefan Arold, and Robert Hoehndorf. 2023. “DeepGO-SE: Protein Function Prediction as Approximate Semantic Entailment.” In Research Square. 10.21203/rs.3.rs-3258432/v1.

Lau, Andy M., Nicola Bordin, Shaun M. Kandathil, et al. 2024. “Exploring Structural Diversity across the Protein Universe with The Encyclopedia of Domains.” Science 386 (6721): eadq4946.

Leclercq, Mickael, and Arnaud Droit. 2026. “Protein Language Models: Applications and Perspectives.” Journal of Proteome Research 25 (2): 507–24.

Lin, Zeming, Halil Akin, Roshan Rao, et al. 2023. “Evolutionary-Scale Prediction of Atomic-Level Protein Structure with a Language Model.” *Science (New York*, N.Y.) 379 (6637): 1123–30.

Liu, Jiachen, Yunxia Chen, Xiuqi Shan, and Huan Wang. 2022. “Association between CILP and IL-1α Polymorphisms and Phenotype-Dependent Intervertebral Disc Degeneration Susceptibility: A Meta-Analysis.” Frontiers in Genetics 13 (October): 1005393.

Liu, Wei, Ziye Wang, Ronghui You, et al. 2024. “PLMSearch: Protein Language Model Powers Accurate and Fast Sequence Search for Remote Homology.” Nature Communications 15 (1): 2775.

Love, Michael I., Wolfgang Huber, and Simon Anders. 2014. “Moderated Estimation of Fold Change and Dispersion for RNA-Seq Data with DESeq2.” Genome Biology 15 (12): 550.

Madani, Ali, Ben Krause, Eric R. Greene, et al. 2023. “Large Language Models Generate Functional Protein Sequences across Diverse Families.” Nature Biotechnology 41 (8): 1099–106.

Moayeri, Mazda, Keivan Rezaei, Maziar Sanjabi, and Soheil Feizi. 2023. “Text-to-Concept (and Back) via Cross-Model Alignment.” In arXiv [Cs.CV]. May 10. arXiv. 10.48550/arXiv.2305.06386.

Özdilek, A. Samet, Ahmet Atakan, Gökhan Özsarı, et al. 2023. “ProFAB-Open Protein Functional Annotation Benchmark.” Briefings in Bioinformatics 24 (2): bbac627.

Pantolini, Lorenzo, Gabriel Studer, Joana Pereira, Janani Durairaj, Gerardo Tauriello, and Torsten Schwede. 2024. “Embedding-Based Alignment: Combining Protein Language Models with Dynamic Programming Alignment to Detect Structural Similarities in the Twilight-Zone.” Bioinformatics (Oxford, England) 40 (1): btad786.

Pedersen, Thomas Lin. 2025. “An Implementation of Grammar of Graphics for Graphs and Networks [R Package Ggraph Version 2.2.2].” Comprehensive R Archive Network (CRAN), August 24. https://CRAN.R-project.org/package=ggraph.

Rembeza, Elzbieta, and Martin K. M. Engqvist. 2021. “Experimental and Computational Investigation of Enzyme Functional Annotations Uncovers Misannotation in the EC 1.1.3.15 Enzyme Class.” PLoS Computational Biology 17 (9): e1009446.

Rives, Alexander, Joshua Meier, Tom Sercu, et al. 2021. “Biological Structure and Function Emerge from Scaling Unsupervised Learning to 250 Million Protein Sequences.” Proceedings of the National Academy of Sciences of the United States of America 118 (15): e2016239118.

Robinson, Mark D., Davis J. McCarthy, and Gordon K. Smyth. 2010. “edgeR: A Bioconductor Package for Differential Expression Analysis of Digital Gene Expression Data.” *Bioinformatics (Oxford*, England) 26 (1): 139–40.

Schnoes, Alexandra M., Shoshana D. Brown, Ivan Dodevski, and Patricia C. Babbitt. 2009. “Annotation Error in Public Databases: Misannotation of Molecular Function in Enzyme Superfamilies.” PLoS Computational Biology 5 (12): e1000605.

Seki, Shoji, Yoshiharu Kawaguchi, Kazuhiro Chiba, et al. 2005. “A Functional SNP in CILP, Encoding Cartilage Intermediate Layer Protein, Is Associated with Susceptibility to Lumbar Disc Disease.” Nature Genetics 37 (6): 607–12.

Sigrist, Christian J. A., Baillie A. Cuche, Estelle de Castro, Emilie Coudert, Nicole Redaschi, and Alan Bridge. 2026. “The PROSITE Database for Protein Families, Domains, and Sites.” Nucleic Acids Research, ahead of print. 10.1093/nar/gkaf1188.

Škunca, Nives, Richard J. Roberts, and Martin Steffen. 2017. “Evaluating Computational Gene Ontology Annotations.” In The Gene Ontology Handbook, edited by Christophe Dessimoz and Nives Škunca, vol. 1446. Methods in Molecular Biology. Humana Press.

Smakowska-Luzan, Elwira, G. Adam Mott, Katarzyna Parys, et al. 2018. “An Extracellular Network of Arabidopsis Leucine-Rich Repeat Receptor Kinases.” Nature 553 (7688): 342–46.

Suzek, Baris E., Yuqi Wang, Hongzhan Huang, Peter B. McGarvey, Cathy H. Wu, and UniProt Consortium. 2015. “UniRef Clusters: A Comprehensive and Scalable Alternative for Improving Sequence Similarity Searches.” *Bioinformatics (Oxford*, England) 31 (6): 926–32.

Synthyra. 2024. “ESMplusplus_large.” Preprint, Hugging Face. 10.57967/HF/3726.

The Gene Ontology Consortium. 2026. “The Gene Ontology Knowledgebase in 2026.” Nucleic Acids Research 54 (D1): D1779–92.

The UniProt Consortium. 2023. “UniProt: The Universal Protein Knowledgebase in 2023.” Nucleic Acids Research 51 (D1): D523–31.

Tipton, K., and S. Boyce. 2000. “History of the Enzyme Nomenclature System.” Bioinformatics (Oxford, England) 16 (1): 34–40.

Wu, Yingcheng, Jiaqiang Ma, Xupeng Yang, et al. 2024. “Neutrophil Profiling Illuminates Anti-Tumor Antigen-Presenting Potency.” Cell 187 (6): 1422–1439.e24.

Xi, Lin, Xu Na Wu, Max Gilbert, and Waltraud X. Schulze. 2019. “Classification and Interactions of LRR Receptors and Co-Receptors within the Arabidopsis Plasma Membrane - an Overview.” Frontiers in Plant Science 10 (April): 472.

Yang, Jianyu, and Shaun Mahony. 2026. “TPCAV: Interpreting Deep Learning Genomics Models via Concept Attribution.” In bioRxivorg. bioRxiv, April 14. 10.64898/2026.01.20.700723.

Yosinski, Jason, Jeff Clune, Yoshua Bengio, and Hod Lipson. 2014. “How Transferable Are Features in Deep Neural Networks?” Preprint. https://arxiv.org/abs/1411.1792.

Yu, Tianhao, Hongzheng Cui, Jinzhi Li, Yunan Luo, Guo-Qiang Jiang, and Huimin Zhao. 2023. “Enzyme Function Prediction Using Contrastive Learning.” Science 379 (6639): 1358–63.

Zanetti, Andrea, Gwendal Dujardin, Lucas Fares-Taie, et al. 2024. “GPATCH11 Variants Cause Mis-Splicing and Early-Onset Retinal Dystrophy with Neurological Impairment.” Nature Communications 15 (1): 10096.

Zeiler, Matthew D., and Rob Fergus. 2013. “Visualizing and Understanding Convolutional Networks.” Preprint. https://arxiv.org/abs/1311.2901.

Zhang, Chengxin, and Lydia Freddolino. 2024. “A Large-Scale Assessment of Sequence Database Search Tools for Homology-Based Protein Function Prediction.” Briefings in Bioinformatics 25 (4): bbae349.

Zhang, Hanyu, Xiting Wang, Chengao Li, Xiang Ao, and Qing He. 2025. “Controlling Large Language Models through Concept Activation Vectors.” In arXiv [Cs.CL]. January 10. arXiv. 10.48550/arXiv.2501.05764.

Zhang, Zhidian, Hannah K. Wayment-Steele, Garyk Brixi, Haobo Wang, Dorothee Kern, and Sergey Ovchinnikov. 2024. “Protein Language Models Learn Evolutionary Statistics of Interacting Sequence Motifs.” Proceedings of the National Academy of Sciences 121 (45): e2406285121.

